# FlowScatt: enabling volume-independent flow cytometry data by decoupling fluorescence from scattering

**DOI:** 10.1101/2020.07.23.217869

**Authors:** Ruud Stoof, Lewis Grozinger, Huseyin Tas, Ángel Goñi-Moreno

## Abstract

**Motivation:** Measuring fluorescence by flow cytometry is fundamental for characterising single-cell performance. While it is known that fluorescence and scattering values tend to positively correlate, the impact of cell volume on fluorescence is typically overlooked. This makes of fluorescence values alone an inaccurate measurement for high-precision characterisations.

**Results:** We developed FlowScatt, an open-source software package that removes volume-dependency in the fluorescence channel. Using FlowScatt, flourescence values are re-calculated based on the unified volume per cell that arises from scattering decomposition.

**Availability:** FlowScatt is openly available as a Python package on https://github.com/rstoof/FlowScatt. Experimental data for validation is available online.

## 1 Introduction

The single-cell characterisations of targeted molecular dynamics via flow cytometry (FC) is at the heart of systems and synthetic biology. From assisting the selection of genetic components to build novel bio-computing circuits (Nielsen *et al*., 2016), to the analysis of complex phenotypes for metabolic engineering applications (Tracy *et al*., 2010), FC is revealed as a fundamental method. Yet, the potential of this technique is often limited by the analysis capabilities of the software package of choice. As a response to this issue, many computational tools are being developed to, among other things, aid with the calibration (Beal *et al*., 2019), stochastic analysis (Tiberi *et al*., 2018) or gating (Becht *et al*., 2019) of FC data.

Gene expression noise (i.e., fluctuations in the concentration of a protein being expressed by a gene) constitutes the *fingerprint* of information processing in a cell, and its characterisation is often the target of FC analysis. The features of noise patterns (e.g., their distribution or signal-to-noise ratios) are fundamental to understand key microbial mechanisms, such as promoter dynamics (Go ñ i-Moreno *et al*., 2017) or infection types (García-Betancur *et al*., 2017). However, noise is not merely a consequence of stochasticity, but also a signal that sometimes (positively) correlates to specific dynamics of the cell. That is the case of cell volume, since the average number of proteins per cell scales with volume (Kempe *et al*., 2015). Therefore, the fluorescence signal of a cell (i.e., the noise of a specific reporter protein), as measured by FC, needs to be calculated based on its corresponding scattering value (FSC), as a proxy for cell volume, for the sake of accuracy. Otherwise, effects of cell volume, an important source of signal variability, would lead to misinterpretations of FC data.

Here we present FlowScatt, a software tool that re-calculates fluorescence information based on scattering values. As a result, the output is presented as volume-independent florescence signals and distributions which can be compared with other experiments.

## 2 FlowScatt

In order to reliably compare FC data between experiments, FlowScatt re-calculates fluorescence distributions for a single value of scattering. To minimize the impact of this process this value is chosen as the mean scattering for the experiments to be compared.

As a first step, the scattering and fluorescence data are log-transformed. The Probability Density Function of this transformed data is estimated and then fit to a bi-variate normal distribution:

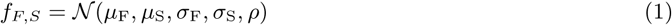

Where *f* is the Probability Density Function described by the binormal distribution 𝒩 and *µ*_x_, *σ*_x_ and *ρ* are the inferred means, variances and covariance of the distribution. *F* and *S* indicate values for fluorescence and scattering, respectively.

Next, the average scattering of all experiments to be compared, 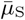 is calculated. Then for each experiment, an estimation of the fluorescence distribution at scattering value 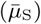 can be obtained using a conditional bivariate log-normal distribution:

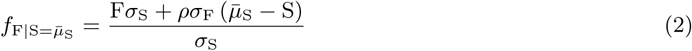

In our example we corrected against the forward scattering height channel to negate volume variation effects, but for small organisms such as bacteria side scattering could be a good measure as well (Felip *et al*., 2007).

## 3 Results

Using FC data as input, FlowScatt uses a filtering process to mitigate the impact of cell volume on fluorescence measurements and adjust their observed distribution appropriately, such that they may be used for comparison across experiments.

We exemplify the utility of FlowScatt by application to FC data from genetic NOT gates in bacterial cells described by Tas *et al*. (2020). Figure 1A shows an individual example experiment which is part of a group of experiments to be compared. The input data (top left) displays a correlation between the chosen fluorescence and scattering channels. If scattering channel distributions vary, for instance due to variation in cell volumes, this correlation will affect comparison between fluorescence distributions of different experiments. This is indeed the case for this example; the binormal distribution fit to this data shows that its mean scattering value (*µ*_S_) lies above that of the group of experiments as a whole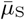. FlowScatt adjusts for this variation to produce a decomposed binormal distribution and filtered data, as shown in Figure 1A (top and bottom right).

**Figure 1:**
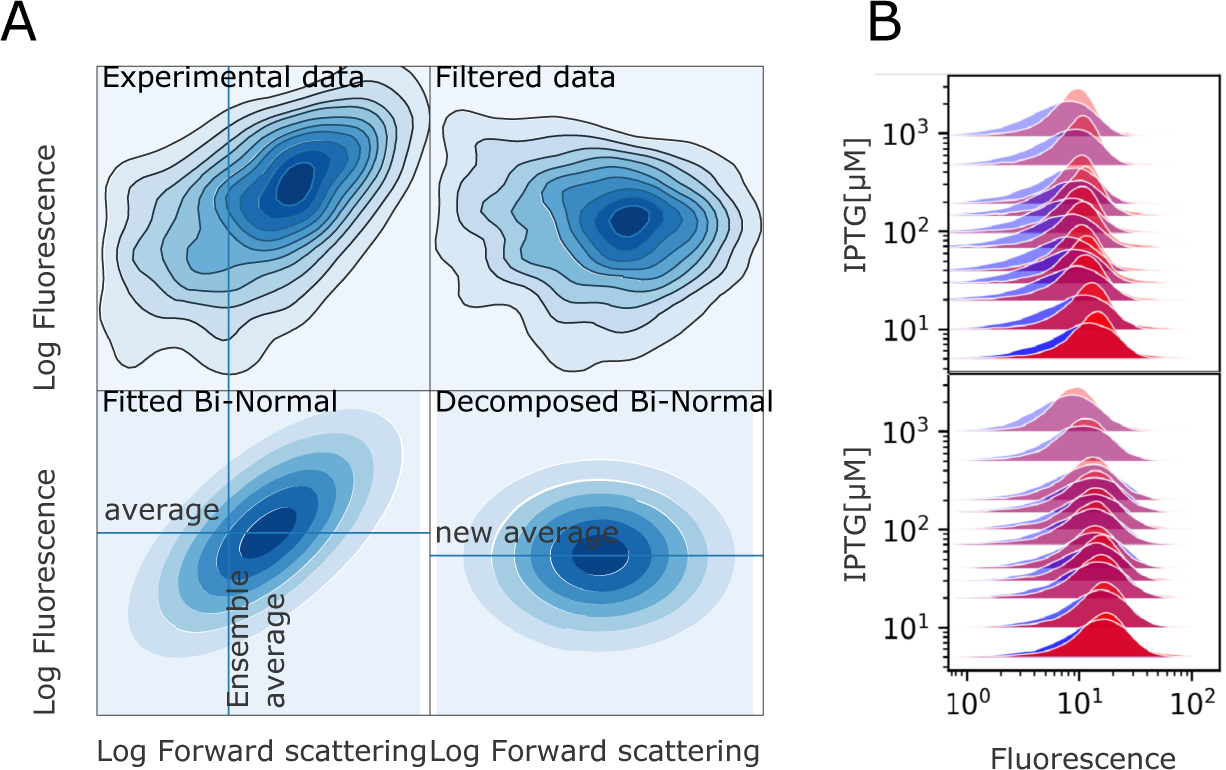
Calculating volume-independent fluorescent data. **A**. Experimental data is fitted to a binormal distribution, then re-calculated based on scattering decomposition (using average) to obtain new, volume-independent, fluorescence data. B. Overlap between experimental (blue) and filtered (red) data. Differences in overlap are due to the top experiments being further below the ensemble average for scattering than the bottom.

In Figure 1 B, differences between observed fluorescence distributions (blue) and adjusted distributions (red) defined by (2) can be seen for two groups of experiments, for which concentration of IPTG is the independent variable. Variability in the extent of overlap between these distributions reflects the variability of the observed scattering distributions for individual experiments.

It is possible that this first iteration of FlowScatt will not completely remove the influence of cell size on fluorescence (if it deviates from a power-law relation). This can be seen when filtered data and the scattering values still correlate (the top right of Figure 1). The tool can be re-run on the new data, however future work could generalise to other experimentally seen distributions.

## 4 Conclusion

Decoupling fluorescence from cell volume produces measurements where signal variability is minimally affected by the growth stage of the microbe. Moreover, results are comparable, not only within the same experiment, but across experiments with varying conditions that may affect the distribution of cell sizes. We advocate the use of FlowScatt as an addition to the efforts directed towards the standardisation of fluorescence measurements.

## Funding

The authors acknowledge the SynBio3D project of the UK Engineering and Physical Sciences Research Council (EP/R019002/1) and the European CSA on biological standardization BIOROBOOST (EU grant number 820699)

## Notes

### Competing Interest Statement

The authors have declared no competing interest.

